# Proteome-wide structural probing of low-abundant protein interactions by crosslinking

**DOI:** 10.1101/867952

**Authors:** Julius Fürsch, Kai-Michael Kammer, Stefan G. Kreft, Martin Beck, Florian Stengel

**Author notes:** These authors contributed equally (Julius Fürsch, Kai-Michael Kammer). Correspondence should be addressed to &.

## Abstract

Proteome-wide crosslinking studies have spurred great interest as they facilitate structural probing of protein interactions in living cells and organisms. However, current studies have a bias for high-abundant proteins. In a paradigm shift, we demonstrate both experimentally and by a kinetic model that this bias is caused by the propensity of crosslinks to preferentially form on high abundant proteins and not by the inability to detect crosslinks due to limitations in current technology. We further show, by using both an in-vitro mimic of a crowded cellular environment and eukaryotic cell lysates, that parameters optimized towards a pseudo 1^st^ order kinetics model result in a significant 3 to 10-fold overall increase in the detection of lower-abundant proteins on a proteome-wide scale. Our study therefore explains the cause of a major limitation in current proteome-wide crosslinking studies and demonstrates how to address a larger part of the proteome by crosslinking.

## Introduction

Proteins and protein complexes are at the center of virtually all biological processes within a cell and deciphering their interactions is key to understand their function. One promising approach for addressing protein-protein interactions is based on the rapidly evolving technology of crosslinking coupled to mass spectrometry (XL-MS). The general approach of XL-MS is to introduce covalent bonds between proximal functional groups of proteins or protein complexes in their native environment by crosslinking reagents. The actual crosslinking sites are subsequently identified by MS and are reflecting the spatial proximity of the respective proteins or subunits in a complex. XL-MS provides a wealth of information on the connectivity, interaction and relative orientation of subunits within a complex, and also contains spatial information in itself, though at relatively low resolution (1, 2).

After some early pioneering work, the field has seen significant technological and conceptual progress over the last couple of years and by now various methods to enrich crosslinks, different crosslinking chemistries, and multiple detection and annotation strategies have been introduced (3, 4). With the structural probing of recombinantly expressed static protein complexes by now being firmly established, the recent application of XL-MS on the proteome-wide level has spurred great interest (5) and an ever-increasing number of studies ranging from bacterial, fungal and mammalian cell lysates and cultured cells (6-9), specific cellular organelles (10-12) and even tissue samples (13) has been reported (for a recent review see (14)). These studies hint at the exciting prospect that XL-MS will soon be able to facilitate the structural probing of interaction partners of any protein of interest within living cells or even organisms.

As exciting as these recent breakthroughs are and even though an impressive number of crosslinks were identified from complex samples, the vast majority of crosslinked peptides that were identified arose known from proteins known to be highly abundant in cells and expressed in large copy numbers, as the ribosome, proteasome, heat shock proteins, histones or enzymes involved in key metabolic pathways as oxidative phosphorylation or the citric acid cycle. We believe that this observation is common to work done independently by various laboratories and raises the question what the cause is of what appears to be a striking imbalance in our ability to detect high abundant or low abundant proteins by proteome-wide XL-MS.

We show, by using both an in-vitro mimic of a crowded cellular environment and eukaryotic cell lysates, that there is indeed a nearly exclusive detection of crosslinks within high abundant proteins in current proteome-wide crosslinking experiments. We demonstrate both experimentally and by a kinetic model that this bias is caused by the propensity of crosslinks to preferentially *form* on high-abundant proteins and as such not by the *inability to detect* crosslinks that have formed on low abundant proteins due to limitations in current MS technology.

We further show that parameter settings optimized towards a pseudo 1^st^ order kinetics model, in particularly an excess of crosslinker, results in a significant increase in the detection of lower abundant proteins within cellular lysates.

Our study therefore not only explains the cause of a major limitation in current proteome-wide crosslinking studies. It also highlights a way forward in how to gain access to a larger part of the proteome for crosslinking studies and paves the way in establishing XL-MS even further as an essential tool for cellular structural biology.

## RESULTS

### Influence of protein abundance on crosslink formation

In order to study the influence of protein abundance on crosslink formation, we mimicked a crowded cellular environment and mixed equal amounts of purified ribosomes from *S. cerevisiae* with increasing amounts of BSA, starting with pure 60S ribosomal particles and subsequently going from a 1 to1 mixture up to a 50-fold excess of BSA (µg/µg), before crosslinking all samples at equimolar concentrations of 1 mM bissulfosuccinimidyl suberate (BS3) (**Figure 1A, Supplementary Dataset 1**).

**Figure 1.**
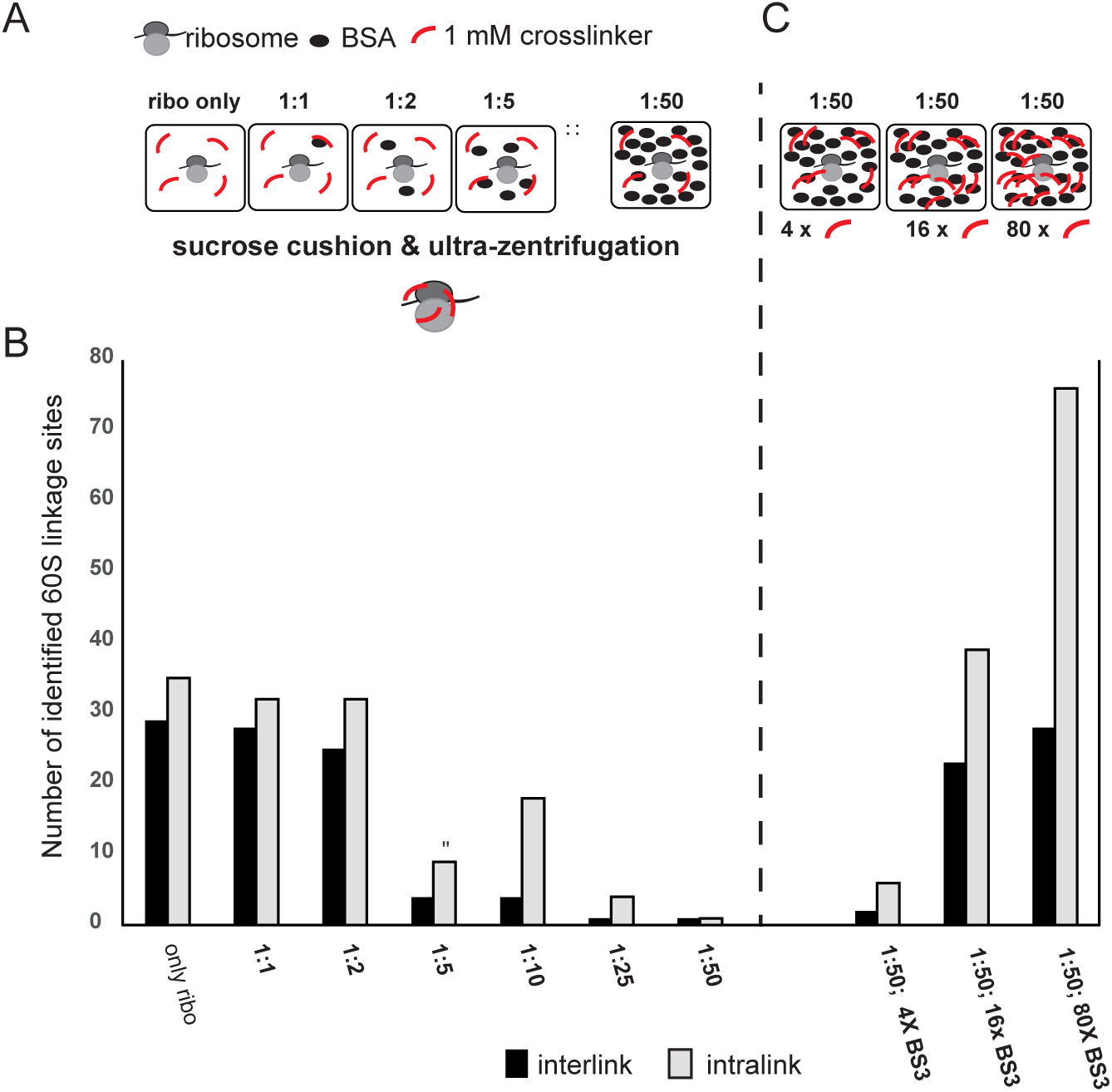
Interdependence of protein abundance and crosslinker concentration on crosslink formation in vitro. (**A**) Equal amounts of purified 60S ribosomal particles from *S. cerevisiae* were mixed with increasing amounts of BSA (ranging from a 1 to1 mixture (µg/µg) up to a 50-fold excess of BSA) and crosslinked at a concentration of 1 mM BS3. After quenching of the reactions, the crosslinked ribosomes were separated again from the BSA by ultracentrifugation using sucrose cushion and prior to analysis by LC MS/MS. (**B**) Detected unique crosslinking sites within the 60S ribosomal subunit (intermolecular and intramolecular) within these samples and (**C**) after addition of excess crosslinker to the 50-fold excess BSA sample - 4x BS3; 16 × BS3; 80 × BS3.

Importantly, after quenching of the reactions, the crosslinked ribosomes were efficiently separated again from excess BSA by ultracentrifugation and prior to analysis by MS (**Figure 1—figure supplement 1**). We chose this set-up, as it enabled us to separate the influence of protein abundance on the *formation* of a crosslink from the possibility that a crosslink has been formed, but could not be *detected* due to current limitations in MS technology (e.g. sensitivity of the MS or chromatographic overlay of a high-abundant with a low abundant peptide).

**Figure 1B** shows the number of detected unique, high-confidence 60S linkage sites (intermolecular- and intramolecular) within the different samples. The data shows a strikingly clear trend where with increasing amounts of additional BSA present, concomitantly less ribosomal crosslinks are formed. This trend is robust and reproducible (**Figure 1—figure supplement 2, Supplementary Dataset 1**).

### Higher crosslinker concentrations increase formation of protein crosslinks in a minimal crowded cellular environment in vitro

To create a challenging scenario, we used 50-fold excess of BSA ratio, where only 3 percent of crosslinks compared to the number using only purified 60S ribosomal particles were still formed, and applied increasing amounts of excess crosslinker (**Figure 1C, Figure 1—figure supplement 2, Supplementary Dataset 1**). Here, a 4-fold increase in the applied crosslinker concentration resulted in a comparable 4-fold increase in the number of formed crosslinks. Increasing the crosslinker concentration by another 4-fold restored the number of formed crosslinks nearly completely (> 95 percent of total crosslinks and 80 percent of interlinks compared to the number formed using only purified 60S ribosomal particles). A further 2-fold increase of crosslinker resulted in an even higher number of formed crosslinks (> 95 percent of interlinks), even though the increase was mainly based on a significantly higher number of intralinks. Our data therefore clearly shows that the addition of access crosslinker significantly increases the number of formed protein crosslinks on our – relatively - low-abundant ribosomal proteins in a concentration-dependent manner and is able to effectively fully reverse the effect that the addition of access BSA protein had on limiting the formation of these ribosomal crosslinks. Taken together they demonstrate the interdependence of protein abundance and crosslinker concentration for crosslink formation in vitro.

### Kinetic model for crosslink formation in lysates

Next, we wanted to test if our findings using a minimal crowded cellular environment in vitro could be reproduced using proteome-wide crosslinking of real cellular lysates. We therefore crosslinked *S. cerevisiae* lysate with disuccinimidyl sulfoxide (DSSO), under conditions (lysate concentration of 1mg/ml and crosslinker concentration of 1mM) described previously as favored for proteome-wide crosslinking (5, 6, 15). To this end we have created a simplified kinetic model assuming equal reactivity for all lysines to describe the reaction kinetics (for a detailed description of our model, including all assumptions and equations, see **Figure 2—figure supplement 1**).

**Figure 2.**
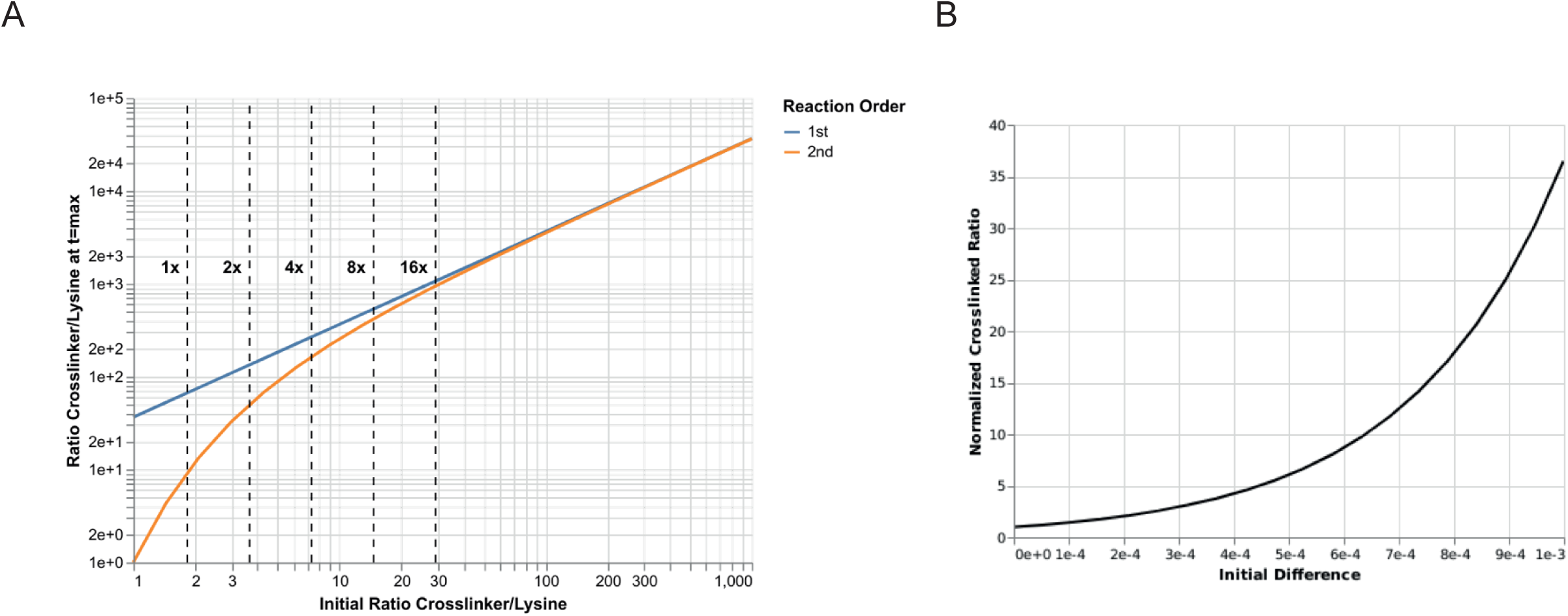
Kinetic model of crosslink formation in lysates. (**A**) Ratio between lysine and crosslinker concentration. The y-axis shows the ratio between crosslinker and lysine concentration at the final time-step where the reaction is supposed to be complete. The x-axis shows the ratio of the initial crosslinker and lysine concentrations. Both axes are in logarithmic scale. The experimental ratios are overlaid with dotted lines. For the low initial crosslinker-to-lysine ratios we assume that the crosslinker concentration is kept constant in the first-order regime. (**B**) Normalized ratio for the crosslinked products versus the absolute initial difference. The y-axis shows the normalized ratio between two crosslinked species at the final time-step where the reaction is supposed to be complete. A ratio of 2 means that compared to the initial ratio the final ratio has doubled. The x-axis shows the absolute difference of the species’ initial concentrations.

Using this model, we find that the previously described lysate crosslinking conditions are almost certainly not sufficient for complete linking of all free lysines, resulting in a suboptimal overall number of formed crosslinks (**Figure 2A**).

Our kinetic model also strongly suggests that high abundant proteins have a higher propensity to react with the crosslinker (**Figure 2B**). The kinetics of the reaction of the crosslinker with the proteins can be approximated with second order kinetics under above conditions. As such the product formation is dependent on both the concentration of the crosslinker and also the concentration of the target protein species. Thus, the likelihood to crosslink less abundant proteins is strongly reduced, or with other words, the high abundant proteins will act like a sponge for the crosslinker. Here we have experimentally tested this concept.

### Proteome-wide crosslinking is limited to high-abundant proteins

We therefore mapped all proteins that were identified by at least one crosslink to a proteome-wide protein abundance distribution in *S. cerevisiae* (https://pax-db.org/species/4932) (16). The protein abundance is here expressed as single protein abundance relative to the total proteome abundance, meaning that a protein with a value of 1000 ppm equals 0.1 % of the total cellular protein amount. **Figure 3A** shows the number of all proteins that were identified by at least one crosslink (inter- or intralink) (blue bars) together with the general distribution of protein abundances in *S. cerevisiae* (orange bars). Overlaid are the cumulative number of proteins (in percent) that were identified at a certain protein abundance for proteins for which a crosslink has been identified (black solid line) versus the complete annotated proteome in *S. cerevisiae* (black dashed line). This data clearly demonstrates that proteome-wide crosslinking is limited to high abundant proteins. We verified this finding by additionally mapping all proteins that were identified in other proteome-wide crosslinking studies spanning different organisms (*E.coli (6)*, mouse (13) and human (5, 6, 9, 15)) and covering lysates (5, 6, 15), cultured cells (5, 6, 9, 15) and specific cellular organelles (13) (**Figure 3—figure supplement 1**).

**Figure 3.**
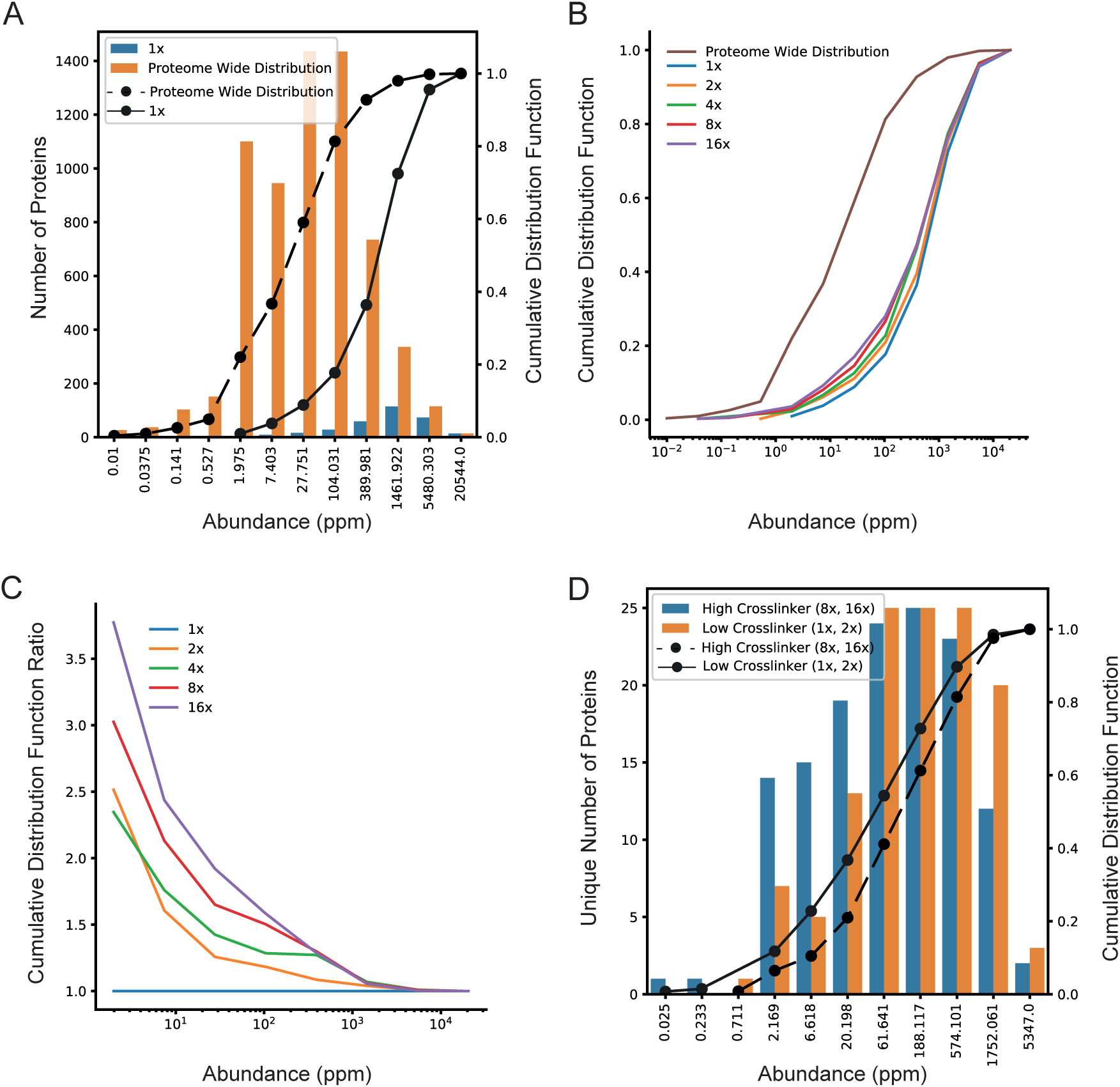
Interdependence of protein abundance and crosslinker concentration on crosslink formation in cell lysates. (**A**) Proteome-wide crosslinking is limited to high abundant proteins. Shown are the number of proteins for which a crosslink (inter- and intralink) could be detected in a proteome-wide crosslinking experiment (blue bars) together with the general distribution of protein abundances in *S. cerevisiae* (https://pax-db.org/species/4932 (16)) (orange bars). The protein abundance is expressed as single protein abundance relative to the total proteome abundance; e.g. a protein with a value of 1000 ppm equals 0.1 % of the cellular protein amount. Overlaid is the cumulative quantity of proteins (in percent) which was identified at a certain protein abundance for proteins for which a crosslink has been identified (black solid line) versus the complete annotated proteome in *S. cerevisiae* (black dashed line). Plotted are the absolute number of identified proteins (y-axis, left) and their cumulative distribution (y-axis, right) versus their respective protein abundance (in ppm) (x-axis). (**B**) Shown are the cumulative distributions of proteins with an identified crosslink using increasing concentrations of DSSO crosslinker (1x (blue); 2x (orange); 4x (green); 8x (red) and 16x (violet) versus the complete annotated proteome in *S. cerevisiae (*brown). (**C**) Depicted are the ratios of these cumulative distributions for these increasing crosslinker concentrations relative to the lowest crosslinker concentration (1mM). The same colour code as in panel B has been applied. (**D**) Shown is the unique subset of proteins for which a crosslink (inter – and intralink) could be detected within samples where a low crosslinker concentration (1x and 2x; blue bars) or a high crosslinker concentration (8x and 16 x; orange bars) was used for crosslinking.

### Proteome-wide interdependence of protein abundance and crosslinker concentration for crosslink formation in cell lysates

Our kinetic model suggests that excess of crosslinker conditions can drive the targeting of lower abundant species. In a next step we therefore wanted to test, if increasing the crosslinker concentration would also increase the number of formed crosslinks on a proteome-wide scale. We therefore crosslinked the same *S. cerevisiae* lysate as above using increasing concentrations of DSSO (1x (blue); 2x (orange); 4x (green); 8x (red) and 16x (violet)) and mapped again all proteins that were identified by at least one crosslink to our proteome-wide protein abundance distribution in *S. cerevisiae* (**Figure 3B, Supplementary Dataset 2**). **Figure 3B** shows an overlay of these cumulative distributions of crosslinked proteins, clearly demonstrating a consistent increase in the number of low-abundant proteins being crosslinked with increasing crosslinker concentration. This trend becomes even more obvious when the relative increase of proteins for which a crosslink was detected within the lysate is compared relative to the lowest concentration of DSSO crosslinker (**Figure 3C**). Here we can clearly see a concentration-dependent effect where the number of detected crosslinks is both dependent on the concentration of the crosslinker and inversely dependent on the concentration of the crosslinked protein. Our data shows a relative increase of detected crosslinked proteins of up to 3.5-fold for low-abundant proteins if the crosslinker is increased by 16-fold compared to the original concentration. In a last step, we took a closer look at the specificity and consistency of the subset of the proteome that can be addressed by XL-MS by varying crosslinker concentrations. **Figure 3D** shows the unique subset of proteins for which a crosslink could be detected in samples when a low crosslinker concentration (1x and 2x; blue bars) or a high crosslinker concentration (8x and 16 x; orange bars) was used. It can clearly be seen that higher concentrations of crosslinker lead to increased crosslinking of lower-abundant proteins, while higher crosslinker concentrations target higher abundant proteins.

**Figure 4** shows respective trends and a similar - but even larger up to 10-fold - increase in the number of crosslinked low abundant proteins using a different crosslinker (BS3) and software suit (*xQuest*), thus corroborating the reproducibility of our results under a different regime (**Supplementary Dataset 3**).

**Figure 4.**
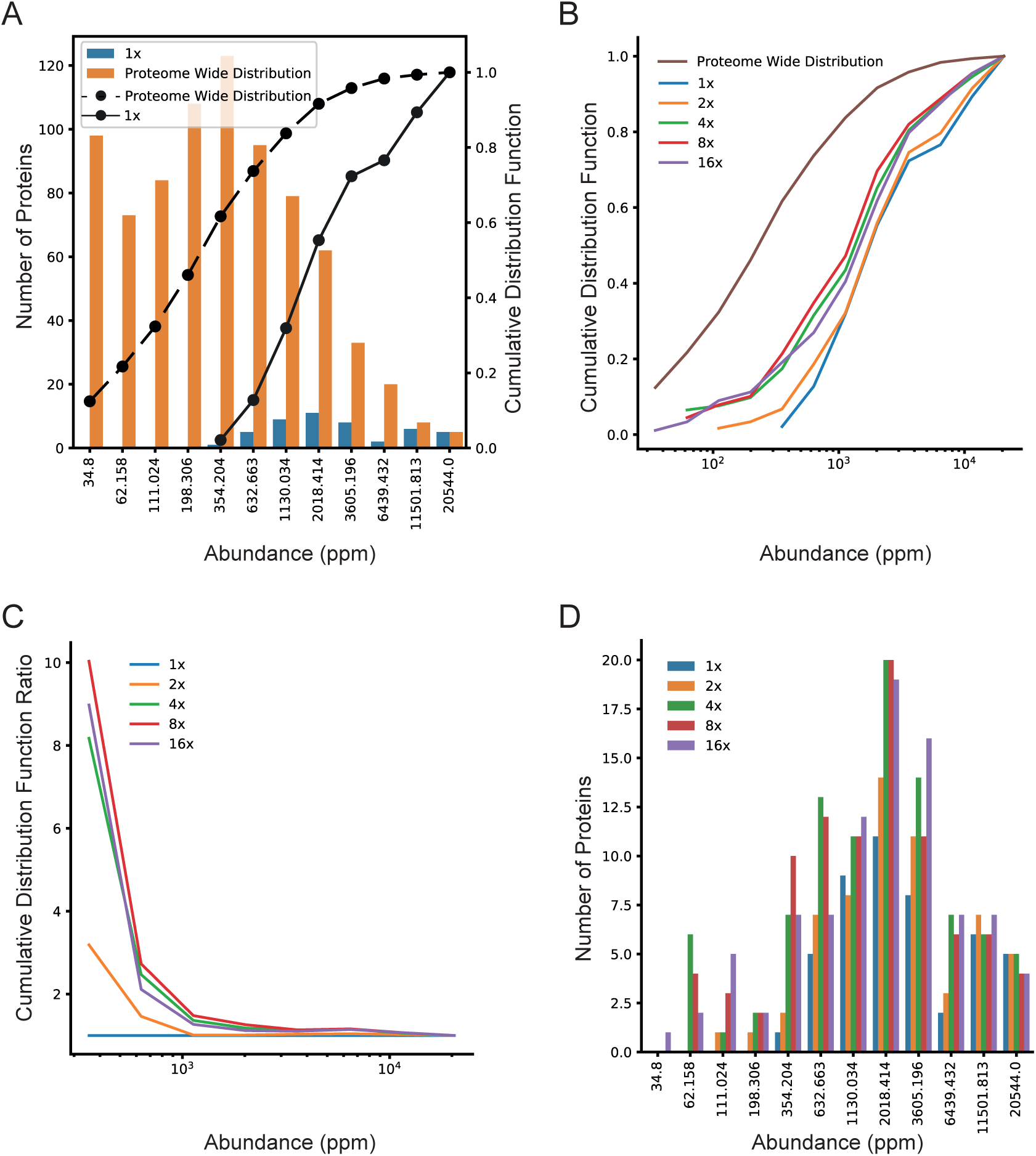
Interdependence of protein abundance and crosslinker concentration on crosslink formation in cell lysates using a different crosslinker and computational pipeline. (**A**) Proteome-wide crosslinking is limited to high abundant proteins. Shown are the number of proteins for which a crosslink (inter- and intralink) could be detected in two proteome-wide replicate experiments using BS3 as a crosslinker (blue bars) together with the pseudo general distribution of protein abundances in *S. cerevisiae* (orange bars). As the non-cleavable linker BS3 could not be searched against the complete proteome, samples were searched against samples were searched against two manually curated databases based on a prior proteome-wide search, one containing 237 proteins (high abundant proteins) and one containing 553 proteins (low abundant proteins) (see methods for a detailed description). For this pseudo proteome-wide distribution only proteins within these databases were plotted. The protein abundance is expressed as single protein abundance relative to the total proteome abundance; e.g. a protein with a value of 1000 ppm equals 0.1 % of the cellular protein amount. Overlaid is the cumulative quantity of all proteins (in percent) for which a crosslink has been identified (black solid line) versus the proteins of the annotated search library in *S. cerevisiae* (black dashed line). Plotted are the absolute number of identified proteins (y-axis, left) and their cumulative distribution (y-axis, right) versus their respective protein abundance (in ppm) (x-axis). (**B**) Shown are the cumulative distributions of proteins with an identified crosslink using increasing concentrations of BS3 crosslinker 1x (blue); 2x (orange); 4x (green); 8x (red) and 16x (violet) versus the proteins of the annotated search library (brown). (**C**) Depicted are the ratios of these cumulative distributions for these increasing crosslinker concentrations relative to the lowest crosslinker concentration (1 mM). (**D**) Shown are the overall number of proteins for which a crosslink (inter- and intralink) could be detected in two proteome-wide replicate experiments using increasing concentrations of BS3 crosslinker. The same colour code as in panel B has been applied.

In summary, these experiments therefore show that proteome-wide crosslinking of cellular lysates leads mainly to the formation of crosslinks on high-abundant proteins but that higher crosslinker concentrations can be used to increase crosslink formation of low-abundant proteins on a proteome-wide level. Taken together they confirm the interdependence of protein abundance and crosslinker concentration for crosslink formation on a proteome-wide scale and show how to gain access to a larger part of the proteome by crosslinking.

## Discussion

This study presents a set of experiments together with a kinetic model where the influence of different factors - as protein abundance and crosslinker concentration – on the formation and detection of (chemical) crosslinks on proteins and protein complexes by MS was studied, both in order to better understand the basic rules that govern this process and to heighten the range and number of proteins that can be addressed in proteome-wide crosslinking studies.

We show that virtually all crosslinks that were identified in a proteome-wide crosslinking experiment are located within proteins or protein complexes that are highly abundant and expressed in large copy numbers, in line with data from previous studies (5, 6, 12, 13, 15). Using an in-vitro mimic of a crowded cellular environment we observe a clear interdependence between the number of detected crosslinks within a given protein complex and the quantity of additional protein present. We then go on to show that this interdependence is caused by the propensity of crosslinks to preferentially *form* on high-abundant proteins.

By using both an in-vitro mimic of a crowded cellular environment and eukaryotic cell lysates in combination with a kinetic model we further show that optimized kinetic parameter settings, in particularly an excess of crosslinker, results in a significant, up to 3 to 10-fold increase in detection of lower abundant proteins on a proteome-wide scale. While we see not only a clear trend towards lower abundant proteins on a proteome-wide scale using optimized parameter settings, we also observe that different crosslinker regimes target a slightly diverse subset of the proteome (**Figure 3D, 4D and Figure 4—figure supplement 1**). This also explains the previously noticed effect of an overall decrease in crosslinked species for higher crosslinker concentrations using SDS-PAGE as a read-out (15). Or in other words, while the previously identified settings of 0.5 to 2 mM of crosslinker concentration for proteome-wide studies (5, 6, 15) are indeed optimal for maximizing the overall amount of crosslinked peptides, we show in this study that this concentration range is not optimal for forming and detecting crosslinks on lower abundant proteins or protein complexes and that an optimized coverage of the proteome will likely only be reached when multiple crosslinker concentration regimes, optimally in conjunction with fractionation of the sample of interest and additional enrichment of crosslinked peptides, are used simultaneously. However, also alternative strategies might be used to address this challenge, such as specific crosslinking of target proteins by the introduction of a chemical handle through genetic code expansion (17).

In summary, our study not only explains a cause of a major limitation in current proteome-wide crosslinking studies. It also demonstrates a way forward how to redesign or repurpose XL-MS studies in the future in order to address an even larger part of the proteome for crosslinking studies and paves the way in establishing XL-MS even further as an essential tool for cellular structural biology.

## Materials and Methods

### Protein expression and purification

Ribosomes were purified from BY4741 yeast cells essentially as described (18). In brief, log phase yeast cells were pelleted, frozen in liquid nitrogen and grounded in a cryo mill (Retsch ®). The lysate was cleared by centrifugation and loaded on a 60% (w/v) sucrose cushion and ribosomes were pelleted by ultracentrifugation. Ribosome pellets were resuspended and incubated with 1 mM puromycin to release nascent chains. Samples were cleared once by centrifugation and loaded on a 10-40% (w/v) sucrose gradient in high salt gradient buffer to split 80S ribosomes into 40S and 60S particles. After centrifugation gradients were fractionated (Teledyne Isco) and 60S particles were collected, concentrated (100,000 MWCO, Merck), and buffer exchanged into buffer A (20 mM HEPES (pH 7.4), 100 mM KCl, 5 mM MgCl2).

### Preparation of *S. cerevisiae* cell lysates

A single-cell colony of *S. cerevisiae* BY4741 was used to inoculate an overnight culture using YPD medium. The main culture was inoculated with an initial OD_600_ of 0.1 and grown at 30°C shaking at 120 rpm. Cells were harvested during exponential growth phase by centrifugation at 4,300 × g for 12 min at 4°C. Cell pellets were washed in cold lysis buffer (50 mM HEPES, pH 7.4, 100 mM KCl, 1.5 mM MgCl2, 0.1 % (v/v) NP-40, 5 % (v/v) glycerol, pefabloc 100 µM, aprotinin and leupeptin 1 mg/L) and centrifuged one more time to get rid of residual medium. Washed cell pellet was reconstituted in 20 mL lysis buffer and quick frozen in liquid nitrogen as small drops. Cell lysis was carried out by grinding the frozen cell droplets in a Retsch® ball mill MM400 at 30 Hz 2 times for 60 s. Frozen cell powder was dissolved in 70 mL lysis buffer on a rolling mixer at 4°C. The cell lysate was cleared by centrifugation at 47,000 × g for 40 min at 4°C. Protein concentration of cleared cell lysate was measured both by BCA and Bradford assay and subsequently diluted to 1 g/L with cold lysis buffer lacking NP-40. The cleared and diluted cell lysate was used for crosslinking experiments.

### Crosslinking of a minimal in-vitro reconstituted crowded cellular environment

Complexes were crosslinked and measured essentially as described (19). In short, equal amounts of roughly 185 µg of purified ribosomes from *S. cerevisiae* were mixed with increasing amounts of BSA using 1:1; 1:2; 1:5; 1:10; 1:25 and a 1: 50-fold excess of BSA (all µg/µg). Samples were subsequently crosslinked at equimolar concentrations (1 mM) of H12/D12 bissulfosuccinimidyl suberate (BS3) (Creative Molecules) for 30 min at 37°C shaking at 650 rpm in a thermomixer (Eppendorf). Alternatively, the 50-fold excess BSA sample was crosslinked with an increasing total amount of BS3 (4x; 16x and 80x) also for 30 min at 37°C (for an exact pipetting scheme see **Supplementary Dataset 4**). After quenching by addition of ammonium bicarbonate to a final concentration of 50 mM, ribosomes were separated from excess BSA by ultra-centrifugation. The crosslinked samples were loaded on a 25 % sucrose cushion and pelleted by ultracentrifugation at 200,000xg for 90 min at 4°C. The complete supernatant was transferred to a fresh tube and stored at −20°C. The ribosome pellet was reconstituted in buffer A by shaking for 1h at 4°C at a vertical shaker. After resolubilization, the ribosome samples were evaporated to dryness, followed by reduction, alkylation, and digestion with trypsin (Promega). Digested peptides were separated from the solution and retained by a solid phase extraction system (SepPak, Waters), and then separated by size exclusion chromatography prior to liquid chromatography (LC)-MS/MS analysis on an Orbitrap Fusion Tribrid mass spectrometer (Thermo Scientific). Protein concentration was normalized both by BCA assay after reconstitution of the ribosome pellet and one more time on the peptide level by A260/A 280 measurement after enzymatic digestion. Data were searched using *xQuest* in ion-tag mode with a precursor mass tolerance of 10 ppm. For matching of fragment ions, tolerances of 0.2 Da for common ions and 0.3 Da for crosslink-ions were applied. Crosslinked samples were prepared in duplicates for all investigated samples, and each of these was measured with technical duplicates. Crosslinks were only considered, if they were identified with deltaS < 0.95 and an Id score ≥ 28.

A list of all identified links can be found in **Supplementary Dataset 1**.

### Proteome-wide crosslinking of lysates

250 μg of 1mg/ml *S. cerevisiae* lysates were cross-linked with increasing total amounts (1x, 2x, 4x, 8x and 16x) of either disuccinimidyl sulfoxide (DSSO) (Thermo Scientific) at 30°C for 1 hour or H12/D12 BS3 (Creative Molecules) for 1 hour at 30°C (for an exact pipetting scheme see Supplementary Dataset 4). After quenching by addition of ammonium bicarbonate to a final concentration of 50 mM, samples were reduced, alkylated, and digested with trypsin (Promega). Digested peptides were separated from the solution and retained by a solid phase extraction system (SepPak, Waters), separated by size exclusion chromatography and after normalization by A260/A280 measurement on the peptide level, subjected to LC-MS/MS analysis on an Orbitrap Fusion Tribrid mass spectrometer (Thermo Scientific).

Samples were measured and searched exactly as described (15). In short, Proteome Discoverer™ 2.2 (Thermo Scientific) with the XlinkX node for crosslinked peptides and SEQUEST HT search engine for unmodified and dead-end-modified peptides was used. Carbamidomethylation (+57.021 Da) was used as a static modification for cysteine and methionine oxidation (+15.996 Da) as variable modification. Data were searched against the Swissprot *S. cerevisiae* database (05/2019) with a 1% FDR cut-off for protein spectral matches.

A list of all identified links can be found in **Supplementary Dataset 2**.

For the BS3 crosslinked samples, measurements and crosslink searches were carried out essentially as described above. Crosslinks were only considered, if they were identified in both replicate samples with a deltaS < 0.95 and with an Id score ≥ 30 in at least one and an Id score ≥ 28 in both replicate samples. As the non-cleavable linker BS3 could not be searched against the complete proteome, samples were searched against two manually curated databases based on a prior proteome-wide search, one containing 237 proteins (high abundant proteins) and one containing 553 proteins (low abundant proteins) (databases were uploaded to PRIDE). All proteins used for creating these libraries were identified throughout all samples by a prior proteome-wide search using Proteome Discoverer™ 2.2. The peptide search was performed using the SEQUEST HT search engine with a precursor mass tolerance of 10 ppm and 0.6 Da for fragment ions. Carbamidomethylation (+57.021 Da) was used as a static modification for cysteine, methionine oxidation (+15.996 Da) and acetylation of the N-terminus (+42.011 Da) as variable modification. Data were searched against the Swissprot *S.cerevisiae* database (05/2019) with a 1% FDR cut-off for protein spectral matches.

A list of all identified links can be found in **Supplementary Dataset 3**.

### Mapping of protein abundances

We used the PAX database (https://pax-db.org/, version 4.1 of the database) to get the relative abundance for all proteins found in the respective species. We used the mapping file available from PAX itself (https://pax-db.org/download, full_uniprot_2_paxdb.04.2015.tsv.zip) to convert the String ID used by the PAX database to Uniprot identifiers. Furthermore, we filtered all proteins not inside the Swissprot database of the respective species (accessed on: 19/06/2019) in order to end up exclusively with curated protein entries. Since the same Uniprot ID can be associated with several String IDs we only kept the most abundant entry in order to end up with a list of unique Uniprot IDs to abundance mappings. These lists were used for reference proteome abundances. Note that for the BS3 experiment we filtered the reference proteins to contain only those that were used for the *xQuest* search. We then assembled a list of unique Uniprot IDs corresponding to the proteins in which we found crosslinks (or the crosslinked proteins from literature) and mapped these to their abundances. The abundance range was discretized into 12 bins with geometric spacing starting from the lowest and going to the highest abundance value found experimentally. Additionally, we added one bin starting at 0, accounting for all reference abundances below the experimental minimum. Since the experimental minimum and maximum abundance are computed for all the different crosslinker concentrations taken together, this sometimes leads to some bins being empty for specific crosslinker concentrations (specifically the low abundance bins for low crosslinker concentrations). The databases were transformed via pandas (version 0.24.2; https://pandas.pydata.org/) and the plots were created using seaborn (version 0.9.0; https://seaborn.pydata.org/) running on Python version 3.7.3.

## Data availability

All data generated or analysed during this study are included in this published article (and its Supplementary information files). The MS data (raw files, databases for *xQuest* search, proteome discoverer and *xQuest* output files) have been deposited to the ProteomeXchange Consortium via the PRIDE (20) partner repository with the dataset identifier PXD014759.

## Acknowledgements

We want to thank Elke Deuerling for purified 60S ribosomal particles. This work was funded by the German Science Foundation Emmy Noether Programme to F.S. (STE 2517/1-1). F.S. also acknowledges funding from the Konstanz Research School Chemical Biology (KoRS-CB) and support from the DFG Collaborative Research Centre (SFB) 969.

## Author Contributions

J.F., K.M.K., M.B., and F.S. conceived the study and experimental approach; S.G.K. purified 60 ribosomal particles from *S. cerevisiae*; J.F. performed experiments and K.M.K. kinetic modelling; J.F., K.M.K., M.B., and F.S. analysed the data, and M.B. and F.S. wrote the paper with input from all authors.

---

**Figure 1—figure supplement 1.**
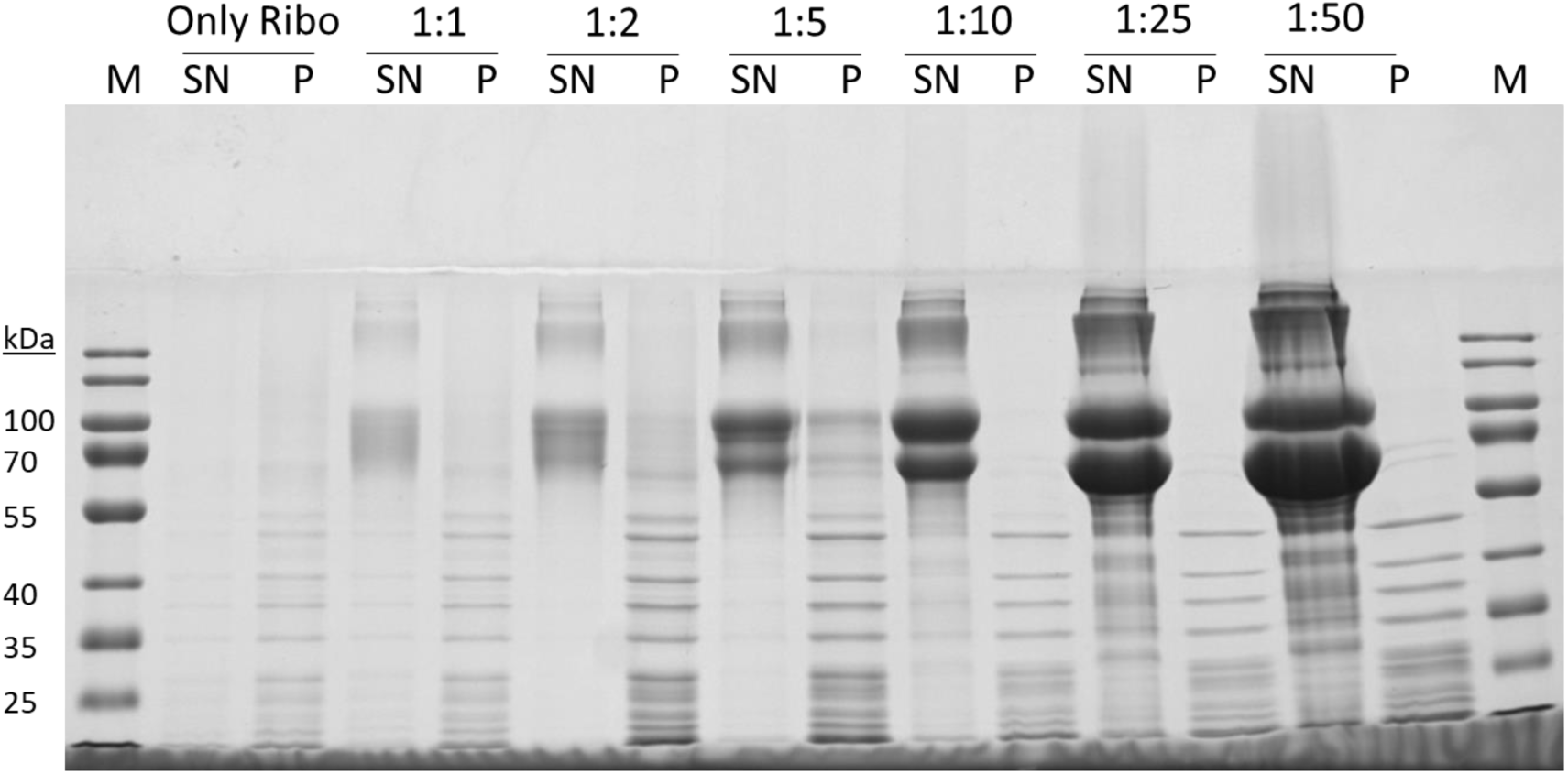
Ribosomes are separated from BSA after crosslinking and prior to MS analysis. SDS-PAGE of supernatant (SN) and pellet (P) fractions of ribosome-BSA mixtures after separation by ultracentrifugation using a 25 % sucrose cushion. Increasing amounts of BSA are seen in the SN fractions, while P fractions show no sign of BSA but clear bands from ribosomal proteins, demonstrating the efficient removal of excess BSA.

**Figure 1—figure supplement 2.**
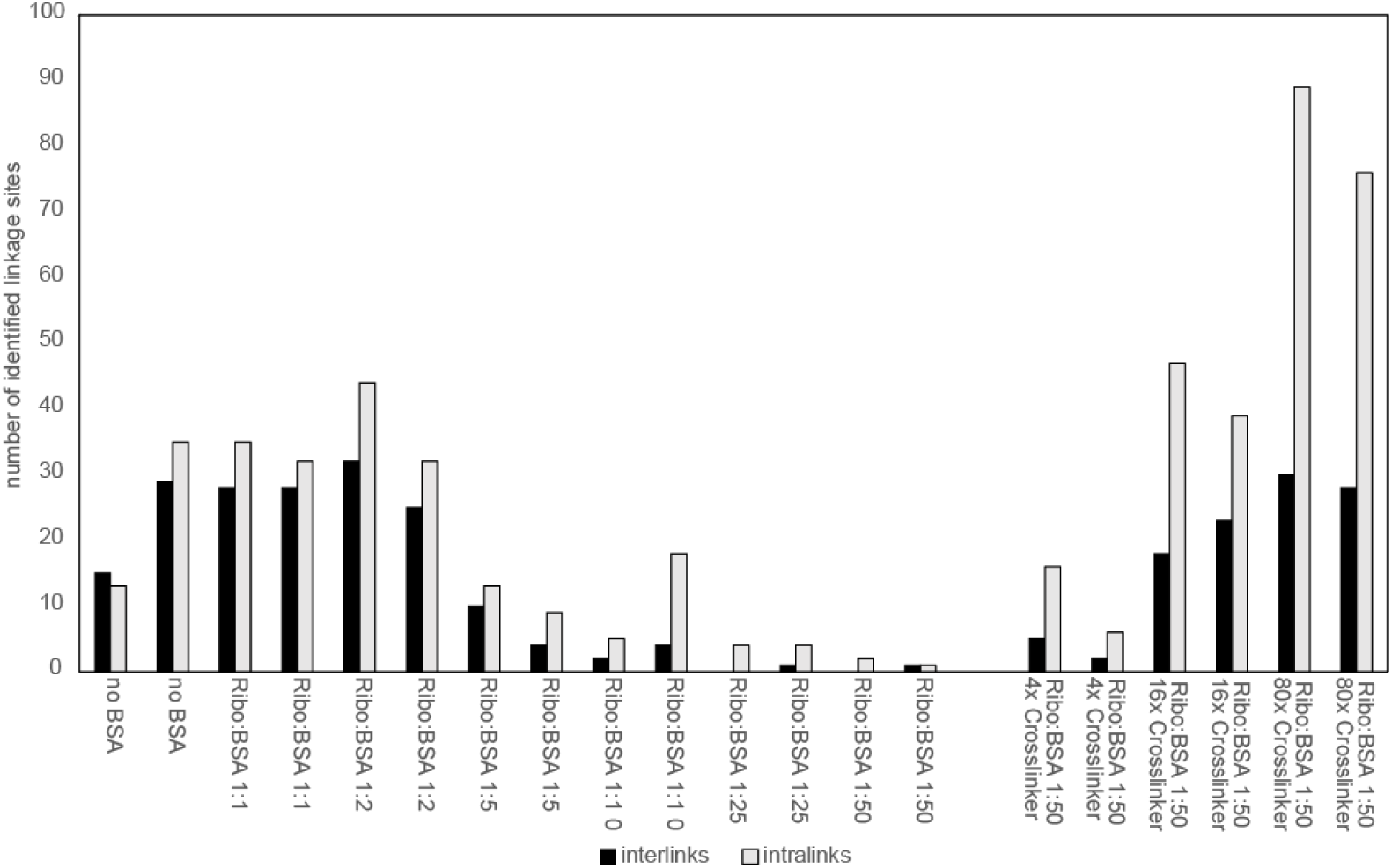
Influence of protein abundance and crosslinker concentration on crosslink formation. Figure 1—figure supplement 2 shows the number of detected unique, high-confidence linkage sites (intermolecular- and intramolecular) within equal amounts of efficiently separated 60S ribosomal particles that had been crosslinked in the presence of increasing amounts of BSA for two replicates. The data shows a very clear and consistent trend, where with increasing amounts of additional BSA protein present, less crosslinks are formed on the 60S ribosome. Upon addition of excess crosslinker, this trend is reversed.

**Figure 2—figure supplement 1. Equations for kinetic model of crosslink formation in lysates**

By establishing a simplified kinetic model, we find that the most common experimental conditions used in literature for lysate crosslinking (lysate concentration of 1 mg/ml; crosslinker concentration of 1 mM, see for example (5, 6, 15)) are not sufficient for completely crosslinking all proteins and favor crosslinking of high-abundant proteins. Looking at a simple model reaction in the form of

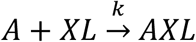

where A represents all lysines found in our mixture and XL the crosslinker. AXL is the (mono)linked product and k is the average rate constant for all lysines. The reaction rate of each species can therefore be described with

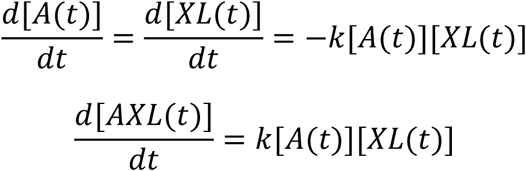

The overall reaction is therefore of second-order with respect to the lysine and crosslinker concentration. When the initial crosslinker concentration [*XL*]_0_ is at least one order of magnitude greater than the overall initial lysine concentration [*A*]_0_, so that [*XL*]_0_ >> [*A*]_0_, the reaction order changes to first-order overall. We assume that

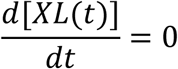

For the free lysine and product reaction rates it follows

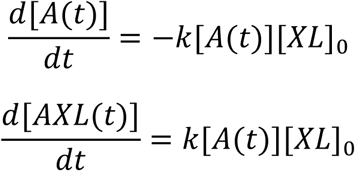

For both the second-order as well as first-order case there are analytical solutions to the differential equations (21). For the second-order case we find

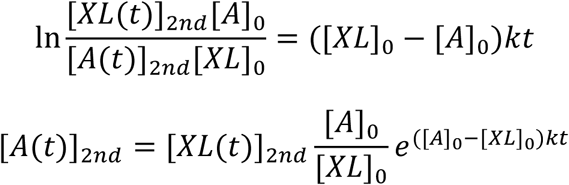

For the first-order case the solution is

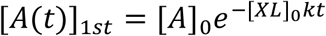

Accordingly, the concentration of crosslinked product is defined by the initial lysine concentration minus the reacted part.

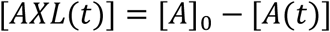

To verify when the first-order-approximation is appropriate we calculate the ratio between the lysine and crosslinker concentration 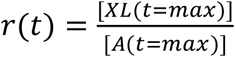 at the end of our reaction where *t* = *max*. We can define this ratio for the first and second order-case:

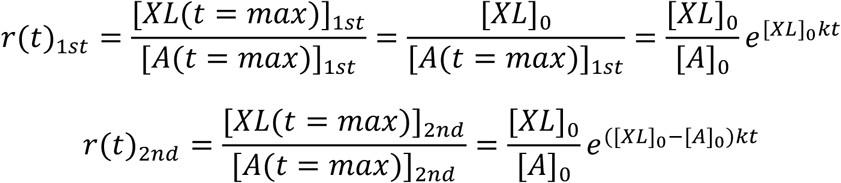

These ratios are plotted in **Figure 2A** below for different initial ratios of crosslinker to lysine. When the two curves approach each other, we are in a good regime for accepting the first-order-approximation. In our experiment the initial protein (and therefore lysine) amount is kept constant while we increase the crosslinker concentration. Our experimental crosslinker factors are overlaid with dotted lines. The parameter values used for the plot are found in the table below.

We approach the first-order-approximation with a crosslinker factor of 8x and greater. When assuming an average lysine frequency of 7% (22) this corresponds to a crosslinker-to-lysine ratio of at least 15. Most crosslinking protocols found in literature are in range of our 1x concentration, corresponding to a crosslinker-to-lysine ratio of ∼2. Note that we always presume our reaction is given enough time for complete turnover of the free lysines which is represented in these models by choosing a large value for *t* = *max* (60 minutes for the experimental lysate crosslinking). The figure should be understood as visualizing a general trend as the actual values for the convergence of the two ratios will strongly depend on the experimental parameters.

In a next step we look at the ratio of two crosslinked species with different abundances for example representing two species inside a cell lysate. For this we define two proteins called B and C (or respectively their free lysines) which both react with our crosslinker. We use the equations of [*A*(*t*)]_1*st*_ and [*A*(*t*)]_2*nd*_ adapted to species B and C to calculate the ratio of crosslinked product for the pseudo-first-order (*r*_*prod*_1*st*_) and second-order (*r*_*prod*_2*nd*_) case:

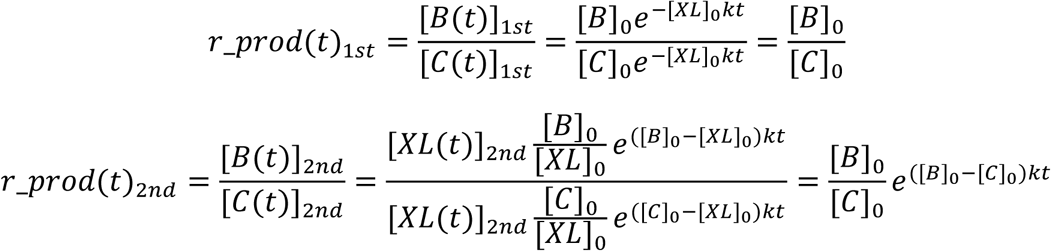

The equation *r*_*prod*_1*st*_ for the pseudo-first-order case demonstrates that under this regime the crosslinked ratio for our different species is just the ratio of the initial concentrations. However, under second-order kinetics the crosslinked ratio also depends on the absolute initial difference between the two species inside an exponential function as well as the rate constant and the time at which the reaction is stopped. To visualize this effect, we normalize *r*_*prod*_2*nd*_ with the initial ratio 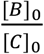:

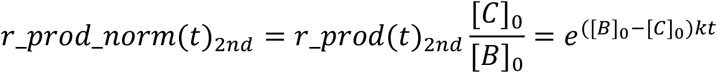

The above equation is plotted in **Figure 2B**. It uses the same parameter values for *k* and *t* = *max* as in the previous figure while the concentration of species B is varied with species C always three orders of magnitude smaller (see table below). The figure shows that in a second order regime we expect a preference for the crosslinker to react with high abundant proteins. It should be taken into account that, as with the previous figure, we just visualize a general trend; the actual magnitude of the effect depends on the values of all the parameters, e.g. *k, t* = *max*, [*B*]_0_, [*C*]_0_.

From these simple models we are able to draw several conclusions. Under first-order conditions the amount of crosslinked protein only depends on the initial free lysine concentration. The initial crosslinker concentration solely influences the speed with which the reaction takes place. In this regime we therefore expect complete crosslinking of all available lysines no matter their respective abundances given enough time. The final ratios between the crosslinked species will be equal to the initial ratios.

Of more interest is the case of second-order kinetics. Here the concentration of our product depends on the instantaneous concentration of both the free lysines as well as the crosslinker. The complete crosslinking of all lysines will take longer since the crosslinker is not as readily available as in the first-order case. Additionally, we now have to consider hydrolysis as well. In aqueous solutions NHS-ester based crosslinkers will readily react with water, decreasing the amount of crosslinker available for reaction (23, 24). This means that for our low crosslinker-to-lysine ratios and for most literature we cannot guarantee complete crosslinking of all available proteins which is especially true for low-abundant proteins since high abundant proteins are linked preferentially. We could therefore lose crosslinks in low abundant proteins where we might be at our detection limit already when their amount is further decreased by incomplete crosslinking. We propose that we can overcome this problem with higher crosslinker concentrations by shifting the reaction kinetics towards pseudo-first-order, thereby reducing the preference for high-abundant proteins.

*Parameter values for plotting the above figures. k was taken from Anjaneyulu* (23) *(for pH 8) while the values for* [*A*]_0_, [*X*]_0_ *and t=max orient themselves at our experimental values.* [*A*]_0_, [*B*]_0_, [*C*]_0_ *were varied within the given range.*

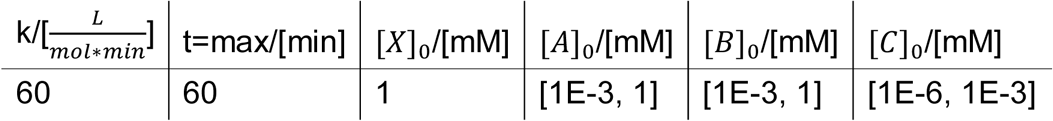

**Figure 3—figure supplement 1.**
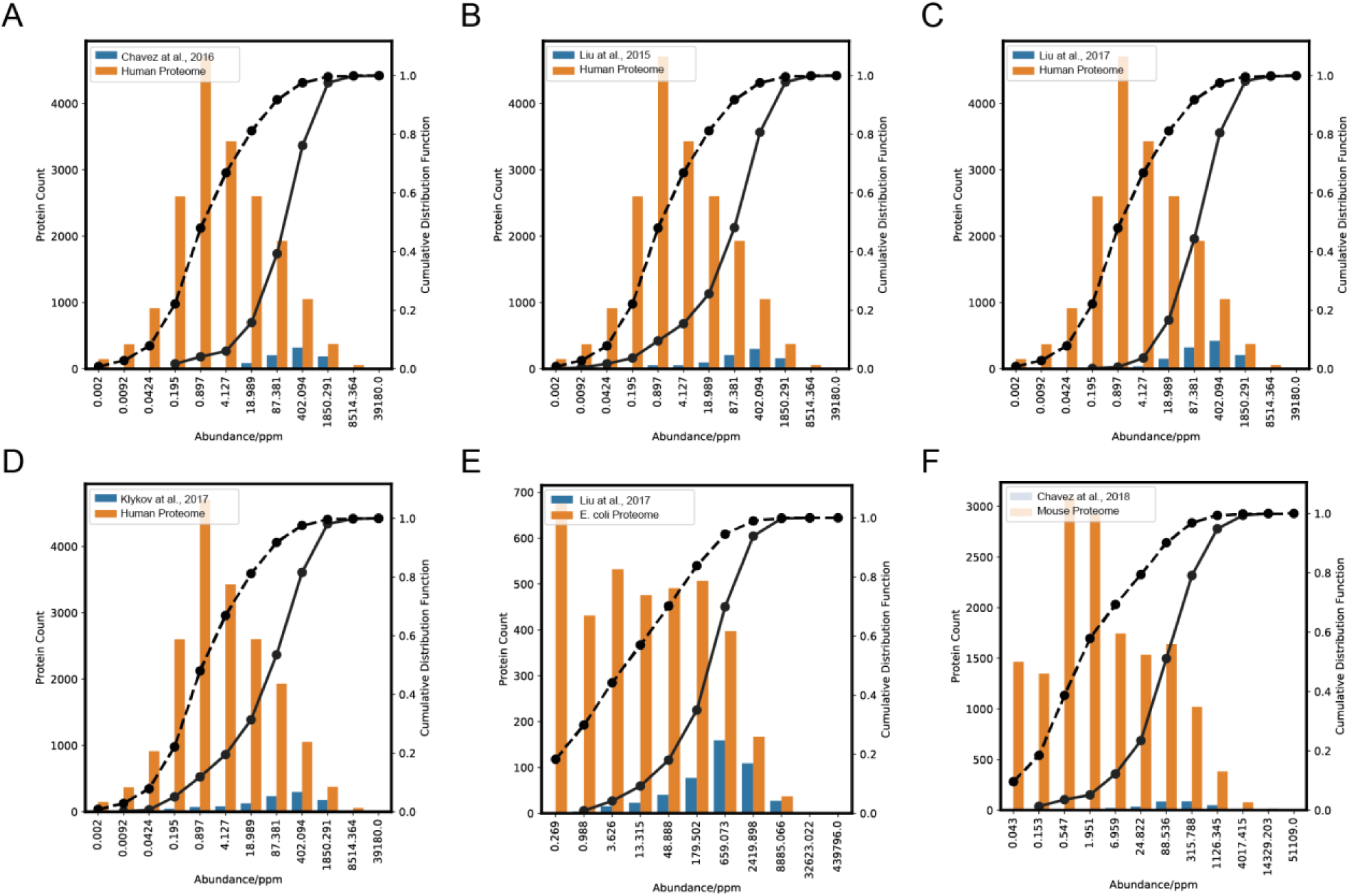
Proteome-wide crosslinking is limited to high abundant proteins. Proteome-wide crosslinking is limited to high abundant proteins. Shown are the number of proteins for which a crosslink (inter- and intralink) could be detected for six different proteome-wide crosslinking experiments (blue bars) together with the respective general distribution of protein abundances in *homo sapiens* (https://pax-db.org/species/9606), *mus musculus* (https://pax-db.org/species/10090) and *E.coli* (https://pax-db.org/species/511145) (orange bars) (16). The protein abundance is expressed as single protein abundance relative to the total proteome abundance; e.g. a protein with a value of 1000 ppm equals 0.1 % of the cellular protein amount. Overlaid is the cumulative quantity of proteins (in percent) which was identified at a certain protein abundance for proteins for which a crosslink has been identified (black solid line) versus the respective complete annotated proteome (black dashed line). Plotted are the absolute number of identified proteins (y-axis, left) and their cumulative distribution (y-axis, right) versus their respective protein abundance (in ppm) (x-axis). Shown are distributions (**A**) Chavez et *al*., 2016 (9), where crosslinking was conducted directly in HeLa cell cultures (**B**) Liu et *al*., 2015 (5), (**C**) Liu et al., 2017 (6) and (**D**) Klykov et *al*. 2019 (15) where crosslinking was conducted in HeLa cell lysates, (**E**) Liu et *al*., 2017 (6), where crosslinking was conducted in *E. coli* cell lysates and (**E**) Chavez et *al*. 2018 (13), where crosslinking was performed in lysates of mouse heart tissue.

**Figure 4—figure supplement 1.**
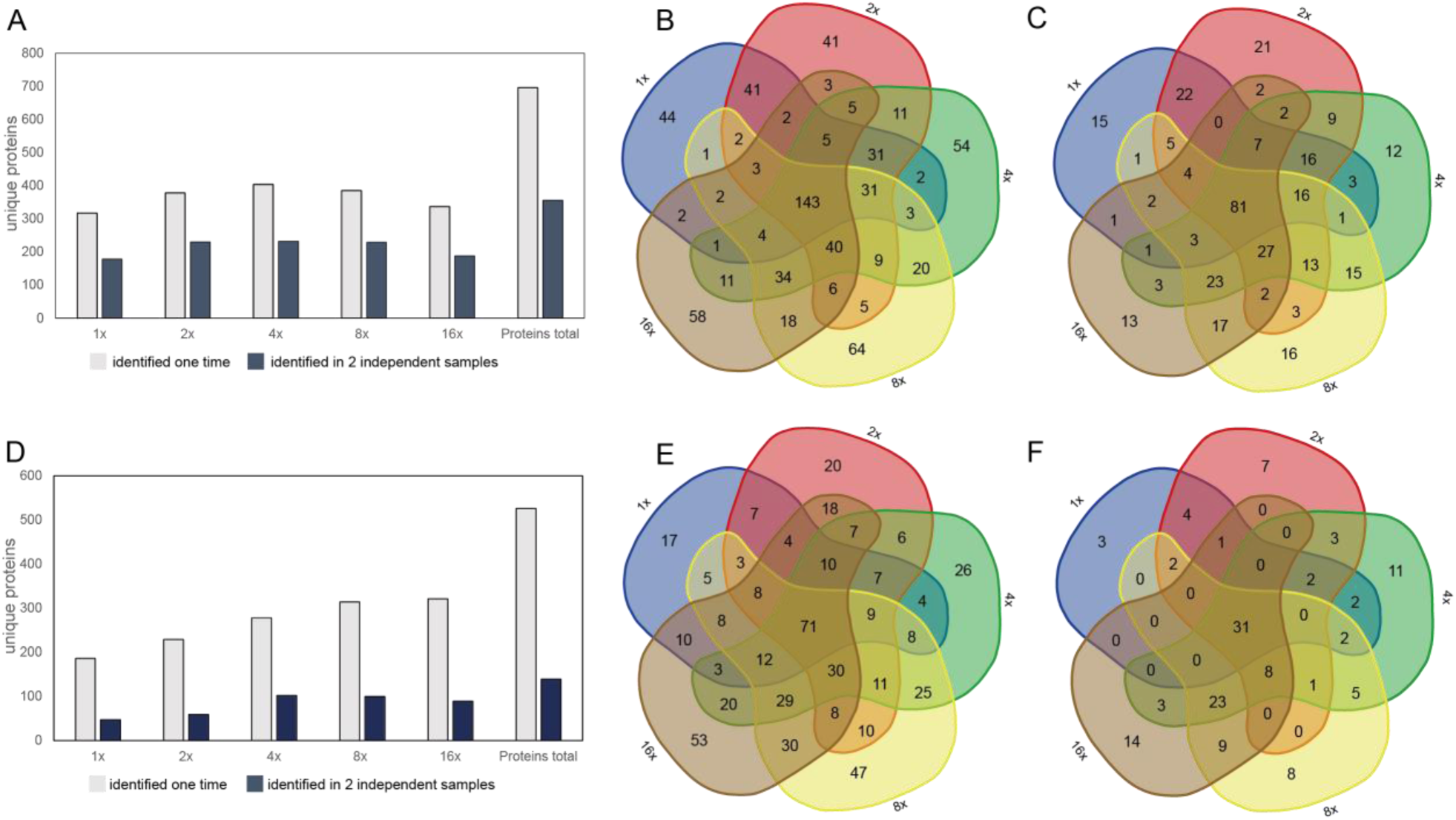
Proteome-wide distribution of identified crosslinked proteins for different crosslinker concentrations. (**A**) Shown is the total number of identified unique proteins for two independent biological replicates in *S. cerevisiae* lysates using increasing concentrations of DSSO crosslinker (1mM (1x); 2x; 4x; 8x and 16x. (**B**) Venn diagram showing the overlap in detected unique proteins between lysates that were crosslinked with different crosslinker concentrations. Here the overlap between neighbouring crosslinker concentrations (e.g. 1x and 2x or 8x and 16 x) is largest (ranging from roughly a third to more than half of all unique proteins identified) and then quite strongly decreases with increasing difference in the used crosslinker concentration (e.g. the overlap between 1x and 16x datasets amounts to only a few percent). The data therefore suggests, that with different crosslinker concentrations, divergent subsets of the proteome can be addressed. (**C**) Venn diagram showing only that subset of proteins, for which crosslinks were identified in both independent biological replicates. (**D to F**) Respective graphs using BS3 as a crosslinker.

**Supplementary Dataset 1: Intra- and inter protein crosslinks of *S. cerevisiae* 60S Ribosome in absence and presence of increasing amounts of BSA or crosslinker**

**Dataset referring to Figure 1 and Figure S1 & Figure S2.**

Shown are the exact amino acid sequence of the crosslinked peptides and the position of the crosslinked lysine residue ‘crosslinked peptide’, the name of the respective protein ‘protein 1’ and ‘protein 2’, nature of the crosslink ‘type’, the position of the crosslinked lysine residues within the UniProt sequence ‘position 1’ and ‘position 2’, the MS raw file in which the crosslink was identified ‘raw file’, ‘delta S’, which gives the delta score of the respective crosslink and is a measure for how close the best assigned hit was scored in regard to the second best, the ‘ld-score’, which is a weighted sum of different scores used to assess the quality of the composite MS2 spectrum as calculated by *xQuest* and the false discovery rate, which was calculated based on the number of decoy crosslinks divided by the number of identified crosslinks in duplicate measurements ‘FDR’.

**Supplementary Dataset 2: Intra- and inter protein crosslinks of yeast BY4741 lysate in presence of increasing amounts DSSO crosslinker**

**Dataset referring to Figure 3 and Figure S4.**

Column labels are the same as described in the legend to Supplementary Data 1.

**Supplementary Dataset 3: Intra- and inter protein crosslinks of yeast BY4741 lysate in presence of increasing amounts BS3 crosslinker**

**Dataset referring to Figure 4 & Figure S4.**

Column labels are the same as described in the legend to Supplementary Data 1.

**Supplementary Dataset 4: Exact pipetting schemes for all in vitro and lysate experiments.**

Data referring to all Figures.

